# The Impact of Digital Histopathology Batch Effect on Deep Learning Model Accuracy and Bias

**DOI:** 10.1101/2020.12.03.410845

**Authors:** Frederick M. Howard, James Dolezal, Sara Kochanny, Jefree Schulte, Heather Chen, Lara Heij, Dezheng Huo, Rita Nanda, Olufunmilayo I. Olopade, Jakob N. Kather, Nicole Cipriani, Robert Grossman, Alexander T. Pearson

## Abstract

The Cancer Genome Atlas (TCGA) is one of the largest biorepositories of digital histology. Deep learning (DL) models have been trained on TCGA to predict numerous features directly from histology, including survival, gene expression patterns, and driver mutations. However, we demonstrate that these features vary substantially across tissue submitting sites in TCGA for over 3,000 patients with six cancer subtypes. Additionally, we show that histologic image differences between submitting sites can easily be identified with DL. This site detection remains possible despite commonly used color normalization and augmentation methods, and we quantify the digital image characteristics constituting this histologic batch effect. As an example, we show that patient ethnicity within the TCGA breast cancer cohort can be inferred from histology due to site-level batch effect, which must be accounted for to ensure equitable application of DL. Batch effect also leads to overoptimistic estimates of model performance, and we propose a quadratic programming method to guide validation that abrogates this bias.

## Main

A standard component of the diagnosis of nearly all human cancers is the histologic examination of hematoxylin and eosin stained tumor biopsy sections. Histologic features identified by pathologists help characterize tumor subtypes, prognosis^1^, and at times can predict response to treatment. Quantification of more subtle pathologic features can further discriminate between good and poor prognosis tumors, such as the quantification of tumor infiltrating lymphocytes in breast cancer, but such detailed analysis can be time consuming and variable between pathologists^2^. The increasing availability of digital histology coupled with advances in artificial intelligence and image recognition has led to computational approaches to rigorously assess pathologic features associated with a variety of tumor specific factors. Deep learning is a subdomain of artificial intelligence, referring to the use of multilayer neural networks to identify increasingly higher-order features of images to allow for accurate classification.

Deep learning on digital histology has exploded as a potential tool to identify standard histologic features such as grade^3,4^, mitosis,^5,6^ and invasion^7,8^. Recently, deep learning approaches have been applied to identify less apparent histologic features, including clinical biomarkers such as breast cancer receptor status^4,9^, microsatellite instability^10,11^, or the presence of pathogenic virus in cancer^12^. These approaches have been further extended to infer more subtle features of disease, including gene expression^13–15^ and pathogenic mutations^16,17^. The predictive accuracy of many of these models have been validated in external datasets, but many studies rely on single data sources for both training and validation.

The Cancer Genome Atlas (TCGA) has been critical for development of deep learning histology models, containing over 20,000 digital slide images from 24 tumor types, along with associated clinical, genomic, and radiomic data^18^. Due to the propensity of machine learning algorithms to overfit, performance is typically reported in a reserved testing set or evaluated with cross-validation, to avoid biased estimates of accuracy^19^. However, the propensity for overfitting of digital histology models to site level characteristics has been incompletely characterized and is infrequently accounted for in internal validation of deep learning models. The genomic batch effects in TCGA and other high throughput sequencing endeavors have been well characterized, and are the product of the hundreds of tissue source sites contributing samples and the multiple sites for genome sequencing and characterization^20–22^. However, histologic imaging data is also prone to batch effect, which may be unique from each tissue submitting site. Prior to sectioning, tissue is first fresh frozen or fixed in formalin and embedded in paraffin, and each fixation method generates unique artifacts^23^. Slides are then stained with the eponymous hematoxylin and eosin stains, the color and intensity of which can vary based on the specific stain formulation and the amount of time each stain is applied. The digitization of slides may then vary due to scanner calibration and choice of resolution and magnification^24^.

Several methods have been proposed to mitigate staining differences between slides^25^, including methods designed for color variation across images by Reinhard and colleaegues^26^, and methods designed specifically for histology by Macenko and colleagues^27^. Color augmentation (Figure 1), where the color channels of images are altered at random during training to prevent a model from learning stain characteristics of a specific site have also been utilized in histology deep learning tasks^28,29^. Most assessments of stain normalization and augmentation techniques have focused on the performance of models in validation sets, rather than true elimination of batch effect^30,31^. Here, we describe the clinical and slide level variability between sites in TCGA, and methods to ensure robust use of internal and external validation to minimize false positive findings with deep learning image analysis.

**Figure 1.**
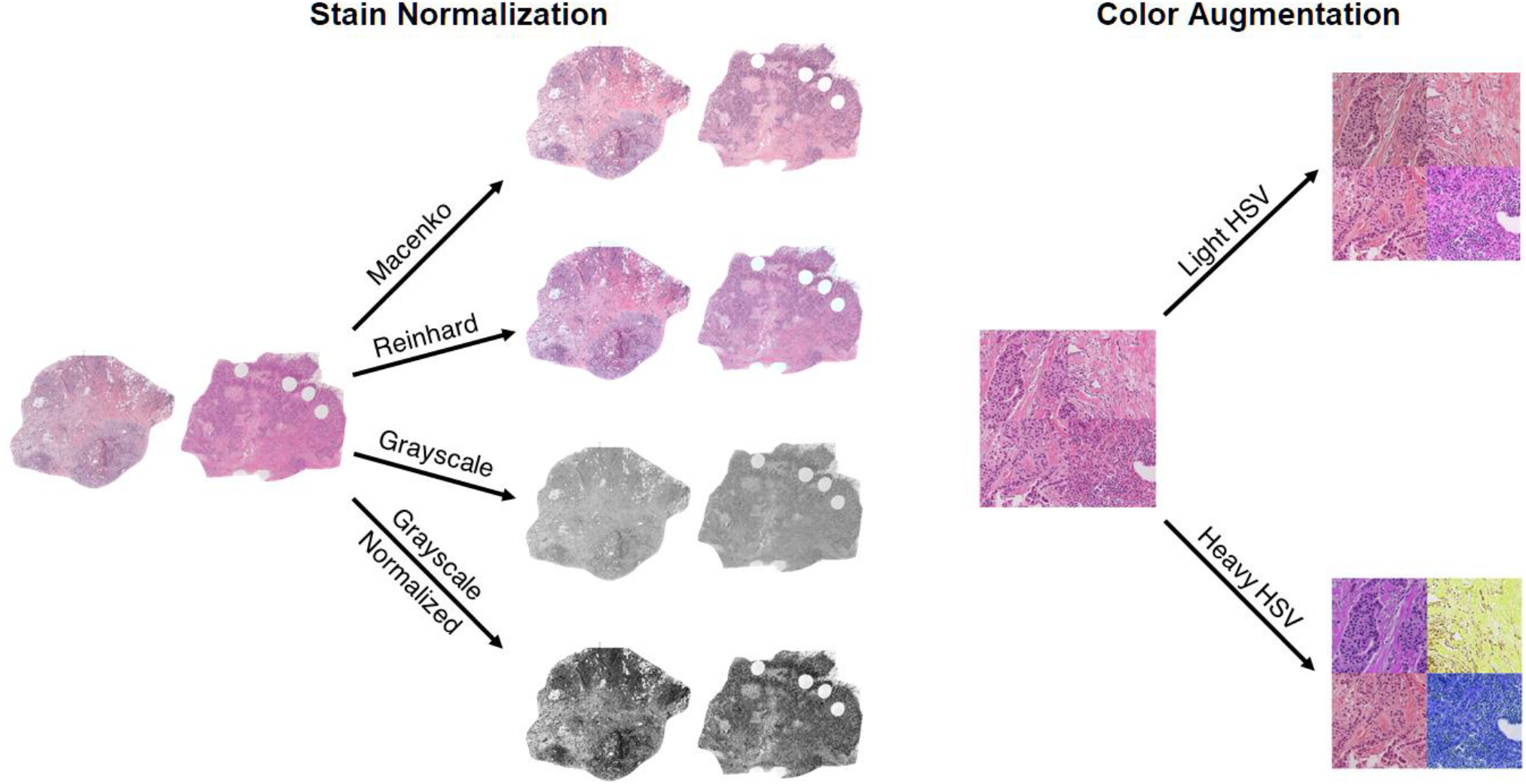
Previously proposed methods of stain normalization and augmentation. Stain normalization refers to changes in color characteristics to reduce the effect of staining differences between slides. Augmentation refers to random variations applied to individual tiles during machine learning to prevent overfitting with regards to the varied characteristic.

## Results

### Characterization of Clinical and Histologic Batch Effect in TCGA

Important clinical variables differ across tissue submitting sites across TCGA. It has been recognized previously that outcomes and survival vary across site for a number of cancers^32^, but even more fundamental factors differ depending on submitting organization. We compared the distribution of basic demographics such as age, race, gender, and body weight index and tumor specific factors such as stage and histologic subtype. Sites were included for comparison if they submitted at least 20 tissue slides. For breast cancer (BRCA TCGA cohort), all demographic characteristics as well as estrogen receptor status (n= 969), progesterone receptor status (n = 969), HER2 expression (n= 847), PAM50 subtype (n = 914), BRCA1 mutational status (n = 931), immune subtype (n = 1,002), and 3-year progression free survival (n = 458)^33^ varied significantly between cohorts, with false discovery rate correction and P < 0.05 (Figure 2). We systematically applied this approach to five other major solid tumor types, and demonstrate that multiple impactful clinical features vary by site for all tumor subtypes tested - including body mass index in colorectal cancer (colon adenocarcinoma (COAD) and rectal adenocarcinoma (READ) TCGA cohorts, n = 476), ALK translocation status in squamous cell lung cancer (LUSC TCGA cohort, n = 157) and lung adenocarcinoma (LUAD TCGA cohort, n = 118), and human papilloma virus (HPV) status in head and neck squamous cell carcinoma (HNSC TCGA cohort, n = 598) – all with P < 0.05 and significant after FDR correction; Supplemental Table 1). Of note given the increasing interest in developing survival models based on pathology, stage varied by site in all cancer subsets tested, and 3 year progression free survival varied across site in all cancers except lung adenocarcinoma.

**Figure 2.**
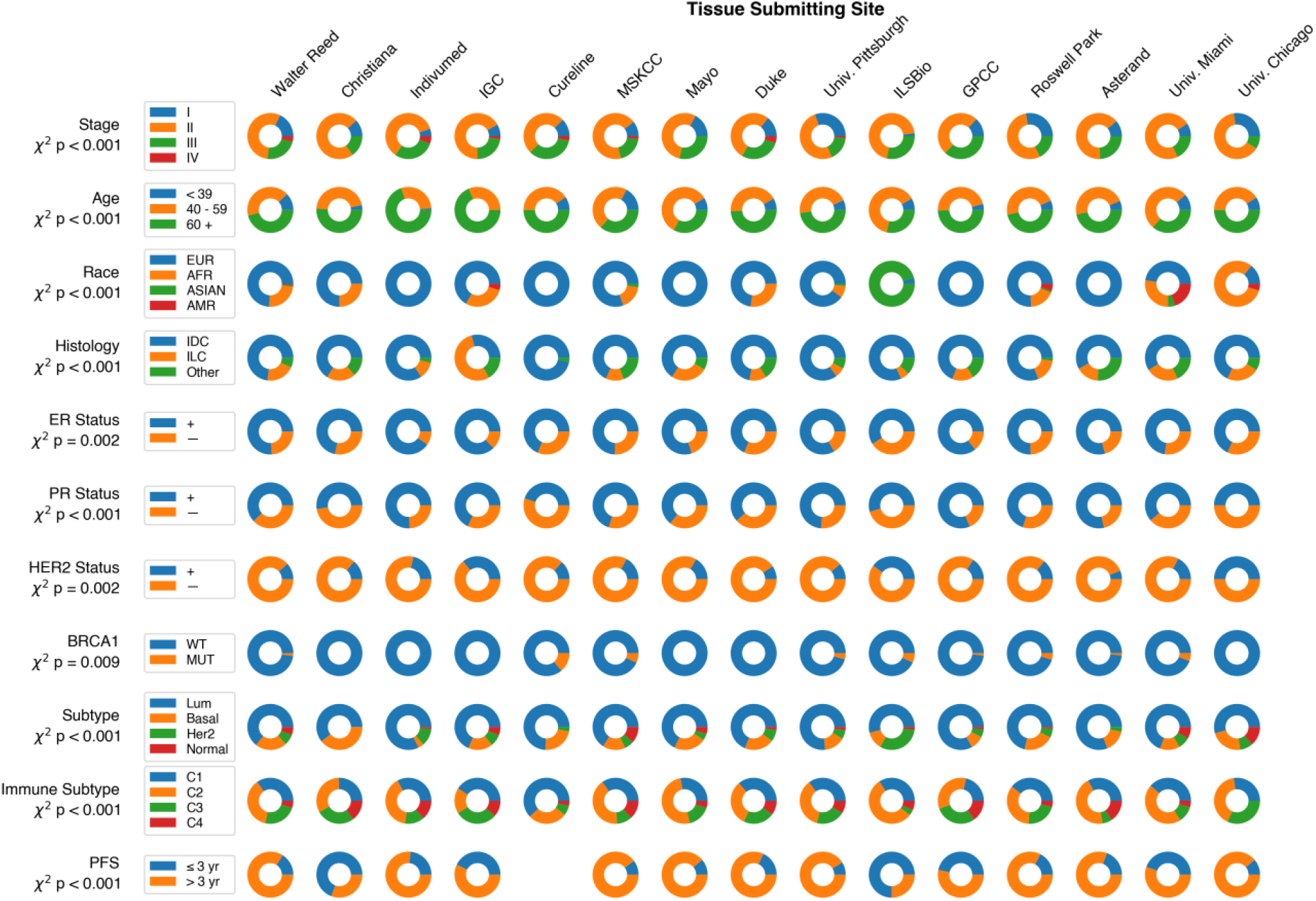
Demographics and Tumor Characteristics of Breast Cancer across Sites with 20 or more samples in TCGA. **Abbreviations: IGC = International Genomics Consortium. MSKCC = Memorial Sloan Kettering Cancer Center. GPCC = Greater Poland Cancer Center. EUR = European, AFR = African, AMR = Native American, IDC = Invasive Ductal Carcinoma, ILC = Invasive Lobular Carcinoma. WT = Wild type. MUT = Mutant. Lum = Luminal. PFS = Progression Free Survival.**

We then applied classical descriptive statistics for image analysis to document the differences in slide features across site, calculating first order statistics and second order Haralick texture features for comparison across sites^34,35^. All first and second order statistics demonstrated variance according to tissue submitting site (Figure 3, Supplemental Table 2). Applying stain normalization techniques at a slide level for breast cancer improved some first order characteristics but measures of dissimilarity for all second order characteristics (as measured by F-statistic) remained greater than that of any first order characteristics (Figure 4). Of note, the second order feature angular second moment remained the most dissimilar feature (highest F-statistic) with any form of stain normalization for all cancer types except lung and head and neck squamous cell carcinoma (Supplemental Table 2).

**Figure 3.**
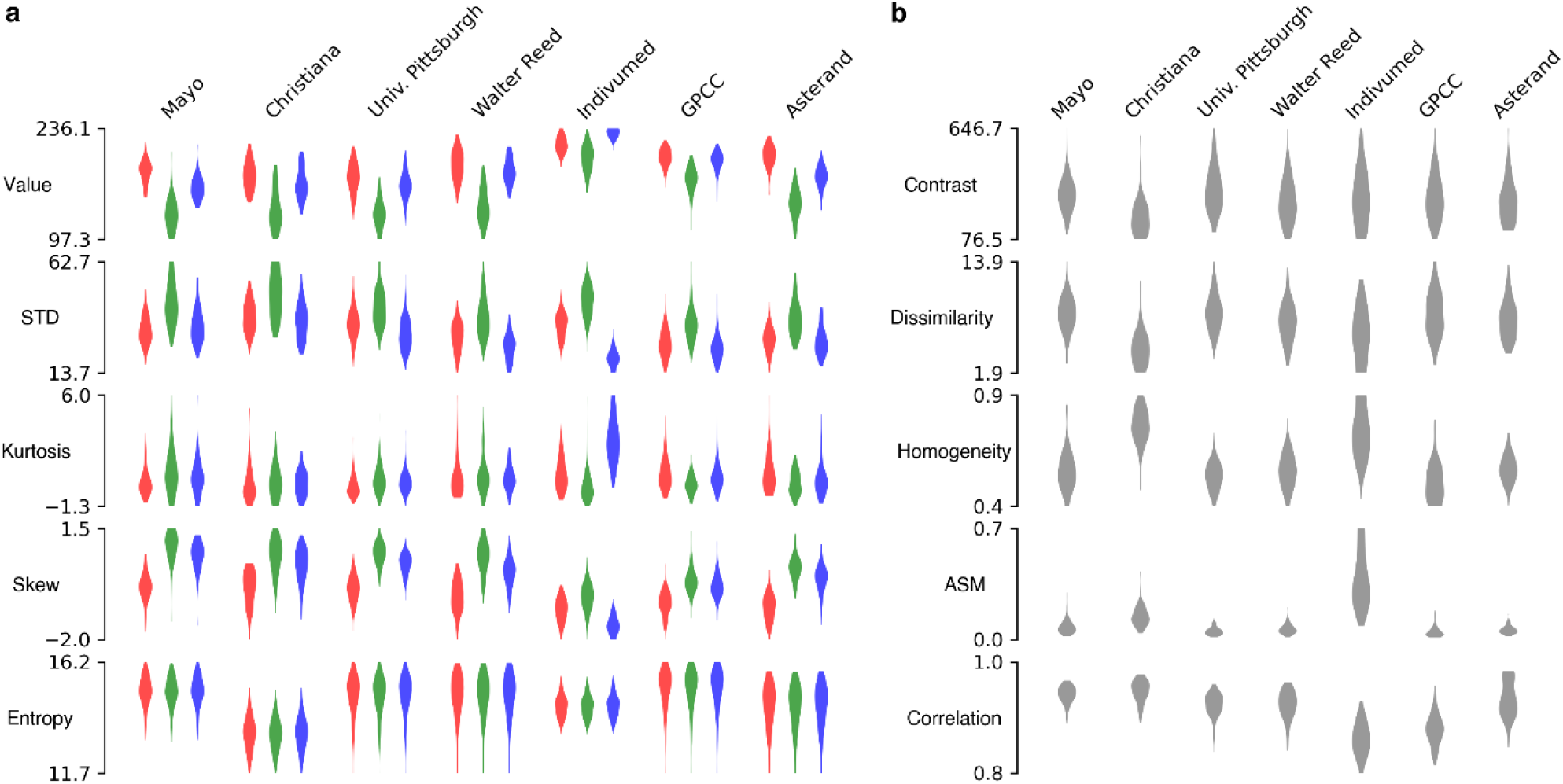
Breast Cancer Digital Histology Feature Variability in TCGA. Sites contributing at least 50 slides are included, demonstrating that image feature variation is not solely a function of small sites that infrequently contributed to TCGA. **a**. First order characteristics for red, green, and blue are shown in their respective colors. **b**. Haralick second order textural features also vary by submitting site. **Abbreviations: STD = Standard Deviation, ASM = Angular Second Moment. GPCC = Greater Poland Cancer Center**.

**Figure 4.**
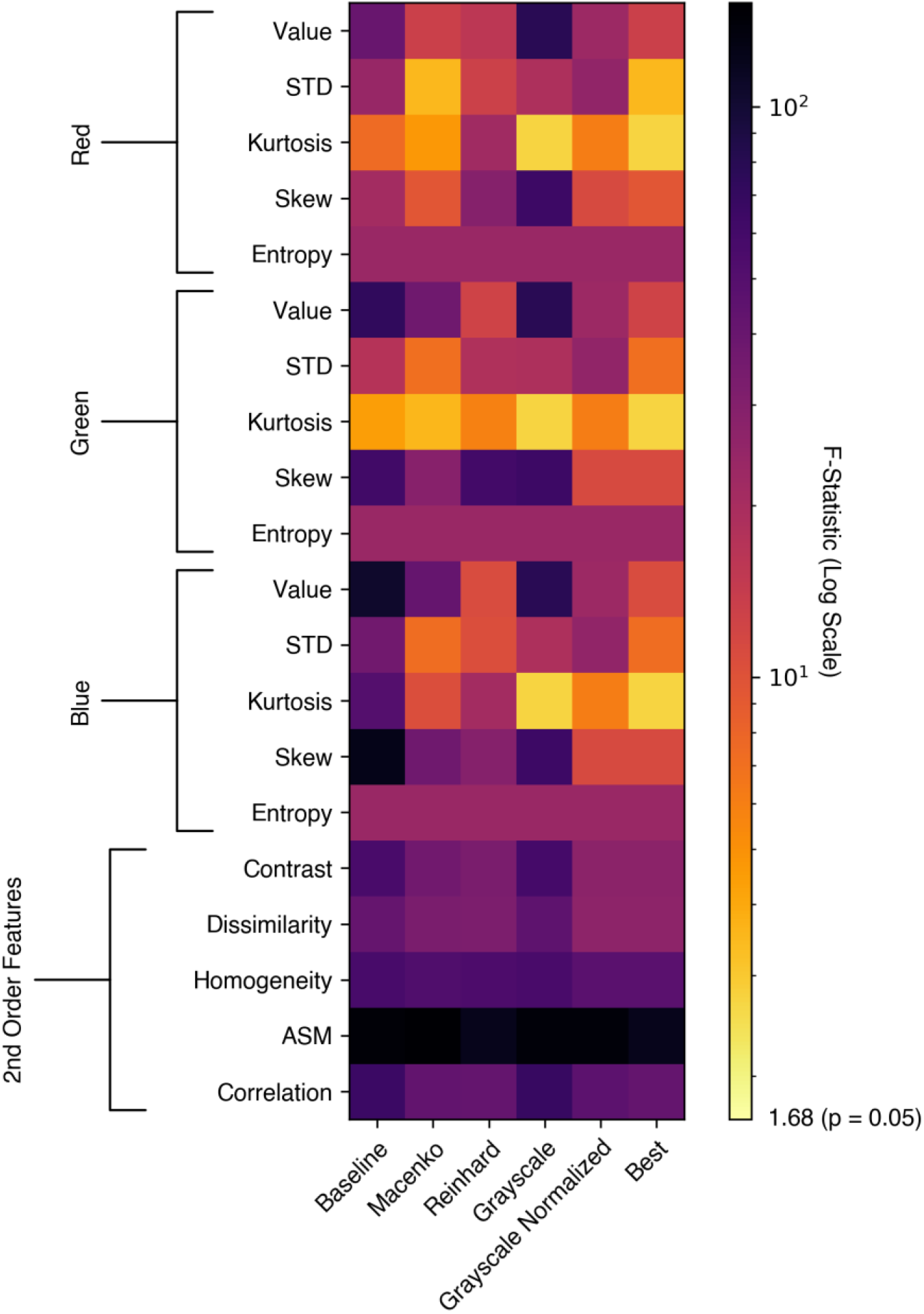
ANOVA F-Statistic for First and Second Order Image Features for Breast Cancer Histology in TCGA. Stain normalization does not resolve first order stain variability (by F-statistic), and minimal impact is seen on second order features. **Abbreviations: STD = Standard Deviation; ASM = Angular Second Moment**

### Deep Learning Algorithms Accurately Identify Tissue Submitting Site

To assess the ability of deep learning to predict tissue submitting site, we trained a deep learning convolutional neural network to predict site. To assess accuracy of site prediction, we used 3-fold cross validation stratified by site (Figure 5a), and calculated the one versus rest area under the receive operating characteristic (AUROC) curve (Supplemental Table 2). The features used by such a model to predict site can be illustrated with a UMAP^36^ representation of final layer activations, with representative slide tiles selected for each UMAP coordinate – in this case demonstrated a clear basophilic-eosinophilic color gradient on the X-axis (Figure 5b). To assess the ability of stain normalization and color augmentation to prevent prediction of site, we repeated this process with normalization or augmentation applied at the tile level for the six examined cancer subtypes (Supplemental Table 3). Site discrimination was highly accurate at baseline, with an average one-versus-rest (OVR) area under the receiver operating characteristic curve (AUROC) ranging from 0.998 for clear cell renal cancer (TCGA-KIRC, n = 513) to 0.966 for TCGA-HNSC (n = 450). Stain normalization techniques modestly decreased accuracy of site prediction, but remained highly accurate with an average OVR AUROC of over 0.850 with all normalization techniques for all cancers. In five of the six cancers, one form of grayscale normalization was able to significantly reduce the site prediction compared to baseline as assessed by paired two-sided t-test.

**Figure 5.**
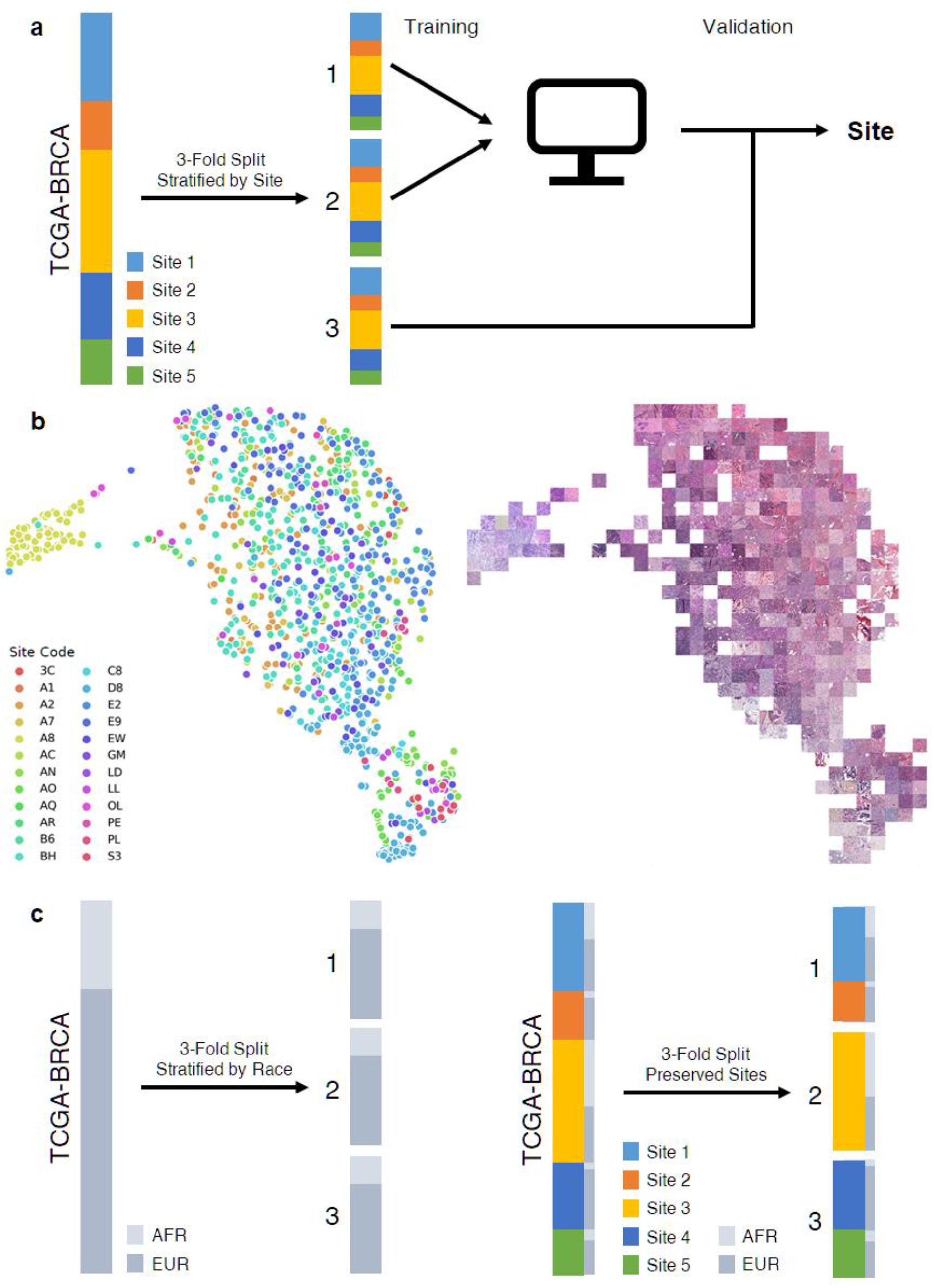
Model Development for Site and Ethnicity Prediction for Breast Cancer Patients in TCGA. **a**. To predict tissue submitting site, data is split into three folds, with each site represented equally in all folds. Cross validation is then performed, where a model is trained on two of the datasets and performance is assessed on the third dataset. This process is repeated threefold for an averaged performance metric. **b**. UMAP representation of final activation of model trained to recognize submitting site. Each point on the left figure represents the centroid tile from a single slide. The nearest tile to each UMAP coordinate is visualized on the right. **c**. We assess the impact of including slides from a tissue submitting site within both the training and validation sets on the prediction of race, using two methods of generating folds for cross validation. First, we split the data into three folds, stratifying by race, irrespective of site. For a comparator, we split the data into three folds where each site is isolated into a single fold, with the secondary objective of equalizing the ratio African and European ancestry in each fold.

Naturally, if a deep learning model can distinguish sites based on non-biologic differences between slide staining patterns and slide acquisition techniques, models will learn the clinical variability between samples at different sites. This is analogous to the Husky versus Wolf problem, where a deep learning model falsely distinguishes pictures of these two canines based on the fact that more wolves are pictured in snow^37^. To demonstrate this fact, we train deep learning models to predict ethnicity, which is highly variable across sites, using two different methods of cross validation (Figure 5c). First, stratifying by race and ignoring site, race can be accurately predicted (Supplemental Table 4). We can correct for biased results by ensuring sites are isolated to single data fold, or **preserved site** cross validation. However, it is still necessary to stratify by the variable of interest, as stratification improves the variance and bias of estimates of accuracy^38^. It is not possible to perfectly stratify by race, but we can select partitions of sites as close as possible for *k*-fold cross-validation using quadratic programming^39^.

This methodology produces perfect stratification (defined as differences by no more than 1 for the number of patients in each category between generated data folds) for preserved site cross validation for all clinical variables described in Figure 2 in the breast cancer dataset, except for gene expression subtype. Notably, estimated AUROC for the prediction of ethnicity in TCGA-BRCA (n = 913) is significantly (p < 0.001 for all comparisons) greater than random chance (which would equate to an AUROC of 0.5) for all stain normalization methods when stratifying by race, but with preserved site cross validation, no model suggests an accuracy greater than random chance (Supplemental Table 4).

We can further demonstrate that models are weighting slide staining characteristics in decision making by examining the predictions for tissue submitting site OL / University of Chicago, the only site where patients of African ancestry comprises the majority of samples. For patients in the validation data folds, false positive predictions for African ancestry (measured at the tile level, n = 2,206) are significantly higher for standard cross validation balanced by race, as compared to preserved site cross validation (Figure 6, Supplemental Table 5). In other words, standard cross validation inaccurately classifies European patients from a site with predominant African ancestry, as the decision is likely related to site level image characteristics. Conversely, the difference in rates of false positive European ancestry is less pronounced, and high rates of false positivity in both models are likely due to the predominance of European patients across training data.

**Figure 6.**
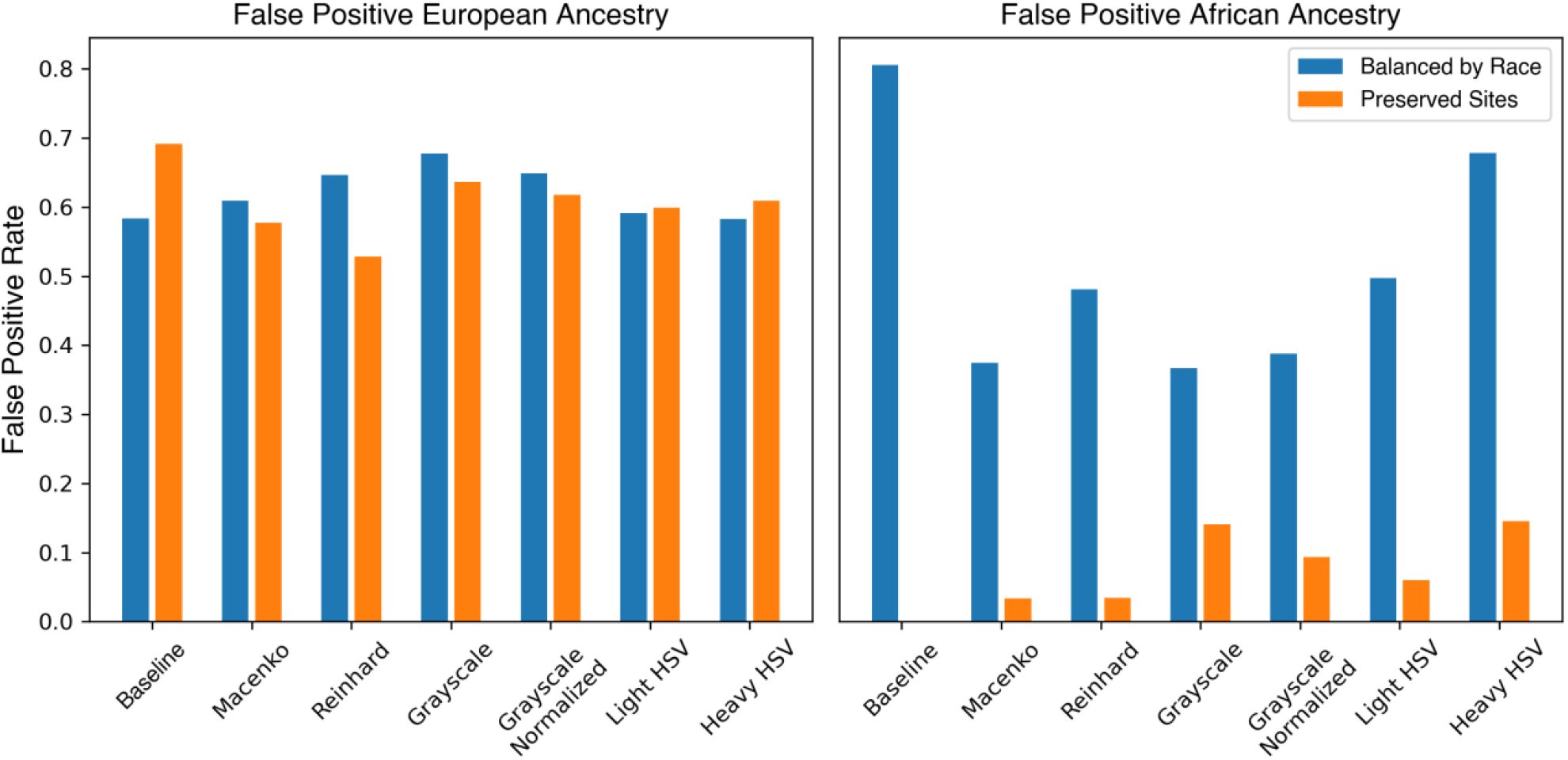
False Positive Rates for Prediction of Ancestry, University of Chicago site. The University of Chicago site submitted a predominance of African ancestry samples to TCGA, and we assess the performance of predictions when models are balanced by race (and University of Chicago patients are thus included in all three folds used for cross-validation), and when sites are isolated to a single fold. False positive rates are similar for European ancestry, but dramatically higher for African ancestry, indicating that models are learning the slide characteristics of samples from University of Chicago and using this information to make calls on patient ethnicity.

## Discussion

We have demonstrated that batch effect exists in the histology images in TCGA across multiple cancer types, and inadequately controlling for this batch effect results in biased estimates of accuracy. Although stain normalization can remove some of the perceptible variation and augmentation can mask differences in color, second order image features are unaffected by these methods, and they do not resolve the ability of deep learning models to accurately identify a tissue submitting site.

Multiple studies have trained and validated models using cases from TCGA without external validation or isolating sites to either the training or validation datasets. Such studies include the classification of cancer histology in lung cancer^17^, genetic mutation prediction in multiple cancer types^16,17,40^, prediction of grade in clear cell renal cancer^41^, prediction of breast cancer molecular subtype^42^, the prediction of gene expression^13^, or correlation of histology and outcome^40,43^. Survival outcomes are particularly challenging to develop rigorous models for using histology from TCGA, and model performance may be falsely elevated not only by the disparate outcomes across sites, but also the site level differences in critical factors relevant to survival such as stage and age. Studies demonstrating histologic discrimination of survival and recurrence in glioblastoma^41,43,44^, renal cell cancer^41^, and lung cancer^45^ patients from TCGA which lack external validation cohorts may have biased estimates of outcome. Prediction of survival may also suffer from this bias^46^ even when correcting for age, stage, and sex, as other factors that vary by site also contribute to outcome, ranging from ethnicity of enrollees, to the treatment available at academic vs community centers. Given that traditional image and textural features vary between sites in TCGA, it is likely that non-deep learning prognostic studies that predict prognosis from traditional image analysis features may suffer from a similar bias^47^. Certainly, there are numerous models which perform well in TCGA without site isolation, and continue to accurately predict outcomes of interest with external validation cohorts, such as prediction of microsatellite instability or BRAF mutations in colon cancer^11,16^. A number of prognostic models have demonstrated the ability of deep learning, trained on a diverse cohort, to accurately predict prognosis in cancers such as colorectal cancer^48,49^ and mesothelioma^50^. However, until the external validity of findings is verified, studies that do not specify the split of sites in their validation and test cohorts must be looked at with a certain degree of scrutiny.

We have also demonstrated that deep learning models have the potential to predict ethnicity from histology *directly* from learned patterns about the demographic makeup of tissue submitting sites. This poses a challenging ethical dilemma for the implementation of deep learning histology models. It has been well documented that black women with breast cancer have a poorer prognosis that is not completely accounted for by stage and receptor subtype^51^. Contributing factors may include delays in treatment initiation and inadequate intensity of therapy^52^. As histology models may be able to identify ethnicity by proxy, they must be carefully implemented in an equitable fashion to avoid inappropriate conclusions^53^. For example, if a prognostic model has learned the staining pattern of a site that recruits primarily African patients with a high rate of recurrence unrelated to disease biology, it may denote low risk patients at that site as ‘poor prognosis’, which could lead to inappropriate administration of chemotherapy or other treatments.

### Best Practices for Addressing Batch Effect for Deep Learning on Histology

We recommend a series of best practices for deep learning studies on histology using TCGA or other combined datasets of multiple hospital sites. First, the variation of outcomes of interest should be reported across included sites. This will allow an assessment of the potential impact histologic batch effect can have on accuracy. Additionally, knowledge about the distribution of outcomes on the training and testing sites can allow for accurate assessment of model performance, as AUROC is an uninformative marker for heavily imbalanced datasets, where the precision recall curve can be more informative^54^. This will also allow an estimate of the impact of batch effect on model predictions. Even if performance stands the test of external validation, models may retain the biases learned from institutional staining patterns. Thus, if outcomes of interest vary heavily across sites, further prospective validation at individual institutions may be necessary before implementation.

If variation of outcomes are seen across sites, models should not be trained and assessed for accuracy on patients from the same contributing site. As we have demonstrated, including sites within the validation and training datasets results in biased estimates of accuracy. The tried and true gold standard for any artificial intelligence endeavor is external validation, which also ensures that not only site level but dataset level batch effects are not reflected in model performance^55^. However, adequate external validation datasets are not frequently unavailable, and it is important to accurately assess the promise of models at an early stage before significant time is spent in further research and investigation. It is rarely possible to perfectly stratify patients by the outcome of interest when isolating sites to either validation or training data, but stratification is still important for unbiased estimates of accuracy. We propose using convex optimization / quadratic programming to identify the split of sites across to allow optimal stratification. This can also be applied to linear outcomes by stratifying the outcome of interest into meaningful subgroups or quartiles prior to optimization.

Finally, stain normalization and color augmentation techniques should still be used to improve model accuracy in external validation and implementation. Although normalization and augmentation do not completely prevent models from learning site specific characteristics, several studies have reported greater validation accuracies with the use of such techniques^30,31^. It is likely that these techniques eliminate some but not all of the reliance that deep learning models have on stain related features; by making the differences in slide characteristics more subtle, models may be more likely to pick up on biologically relevant factors. Grayscale normalization likely discards some relevant biologic information^56^, but the ideal method for normalization has not yet been definitively determined, and may vary per application of interest. Similarly, although attempting to normalize second order features derived from the gray level co-occurrence matrix may render sites more indistinguishable, such features are closely associated with intrinsic tumor biology and must likely be preserved for deep learning applications^57^.

### Limitations

We present a comprehensive description of pixel level characteristics across site in TCGA using classical image analysis techniques, however other factors likely contribute to the detectable differences between sites. Although the compression factor and resolution also varies between the slide files obtained at different sites, we did not assess the ability of deep learning models to detect these factors. It is likely that other higher order image features contribute to the site level differences, such as Gabor, wavelet packet, and multiwavelet features.^24^ However, extensive characterization of all described textural features is not necessary to demonstrate the presence of batch effect and the impact this has on model performance.

We have chosen a subset of proposed stain correction methods, but there have been innumerable approaches and variations that may further reduce the intra-site variability in staining. An unsupervised learning approach to normalizing stains has been proposed, but did not outperform methods of augmentation without normalization in test datasets^30^. Adversarial networks may also allow for models to avoid learning undesirable characteristics of datasets^58^.

## Conclusions

Digital histology in TCGA carries a multifactorial signature that is characteristic of the tissue submitting site. This signature can be easily identified by deep learning models, and can lead to an overestimation of model accuracy, if hospital sites are included in both the training and validation datasets. Care should be taken to describe the batch effect of outcomes of interest across sites, and if significant, a submitting site should be isolated to either the cohort used for training or for testing a model. A quadratic programming approach can maintain optimal stratification while still isolating submitting sites to either training or validation datasets.

## Methods

### Patient cohorts

Patient data and whole slide images were selected from 6 of the tumor types from TCGA with the highest number of slides available to better identify site to site batch effect. Tumor types included breast (BRCA)^59^, colorectal (COAD and READ – with data combined for sites enrolling to both cohorts)^60^, lung squamous cell carcinoma (LUSC)^61^, lung adenocarcinoma (LUAD)^62^, renal clear cell (KIRC)^63^, and head and neck squamous cell carcinoma (HNSC)^64^. Slides and associated clinical data was accessed through the Genomic Data Commons Portal (https://portal.gdc.cancer.gov/). Ancestry was determined using genomic ancestry calls as per Carrot-Zhang and colleagues^65^. Immune subtypes were used from the work published by Thorsson and colleagues^33^. Informed consent was obtained for all participants in TCGA.

### Image Processing and Deep Learning Model

Scanned whole slide images of hematoxylin and eosin stained tissue were acquired in SVS format. Each slide was reviewed by a pathologist for manual annotation of area of tumor, to ensure ink or other non-cancer artifacts did not influence slide level statistics^31^. For slide level first order and second order statistical analysis, slides were downsampled to 5 microns per pixel, or approximately 2X magnification. For deep learning applications, the tumor region of interest is tessalated into 299 pixel x 299 pixel tiles for evaluation, each representing a 302 μm x 302 μm area of histology, effectively generating an optical magnification of 10X. Convolutional neural network models are written in Python 3.8 using TensorFlow and structured based on Xception^66^ with pretraining weights used from ImageNet^67^. This is followed by a single hidden layer with width 500 and a softmax layer for prediction. Further details regarding implementation have been previously published^16,19^.

Each tile is assigned a label associated with the outcome of interest. Tile libraries were also balanced by category to eliminate bias, such that the number of tiles for each target category was equivalent. Models are trained to categorical outcomes using the Adam optimizer and sparse categorical crossentropy loss. Stain normalization and augmentation is applied to individual tiles at the time of training and assessment. Macenko and Reinhard normalization is applied as previously described^26,27^, grayscale refers to direct slide conversion to grayscale, and ‘grayscale normalized’ refers to conversion to grayscale with histogram equalization^68^. Both light and heavy levels of hue saturation value (HSV) augmentation was applied, with light augmentation multiplying each of these three channels by a scalar from 0.9 to 1.1, and heavy augmentation multiplying the hue and saturation channels by a random scalar from 0.7 to 1.3. Additionally, further augmentation through random tile rotation is performed, and further normalization ensures inputs have a mean of zero and variance of one. Models are trained with 3-fold cross validation, learning from two splits of the data and then evaluated on the third split (Figure 5). Models were trained on 3 epochs of the training dataset. Deep learning model training and evaluation was performed on 16 deep learning-specific NVidia Tesla V100s graphical processing unit (GPU) nodes within a HIPAA-compliant environment.

### Statistics

To quantify differences between categorical clinical features across sites, a Chi-squared test is used, with significance determined using a false discovery rate (FDR) of 0.05 with the Benjamini-Hochberg method. A similar approach is applied using ANOVA to identify differences between slide level characteristics between sites, with the same FDR. First order statistics are calculated from individual red, green, and blue pixel values across images, and include mean, standard deviation, skewness, kurtosis, and entropy, the latter being calculated as follows:

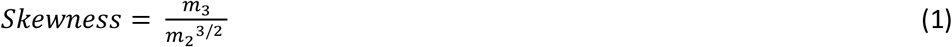

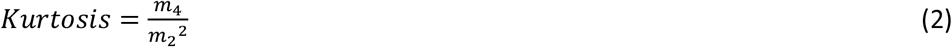

Where 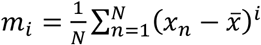

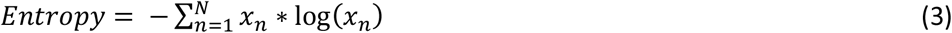

Second order Haralick features^35^ were calculated from the gray level co-occurrence matrix *P*:

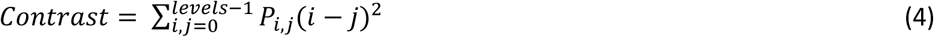

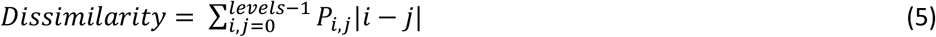

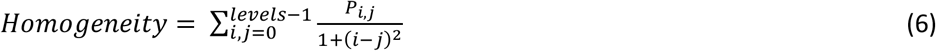

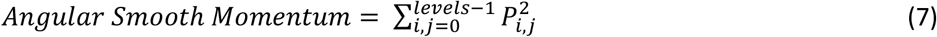

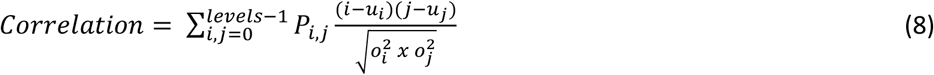

Similar values for calculated features were seen for angles of 0°, 45°, 90°, and 135° so reported values for second order features are averaged across these four angles. The AUROC values are reported as means and ranges generated with 3-fold cross validation. Comparisons between average AUROC for prediction of site with different methods of stain normalization was performed with a two sided paired t-test with an FDR of 0.05. To calculate *k*-folds for cross-validation where each site is isolated to a single fold, we define the following convex optimization problem. If *m*_*s,c*_ is a binary variable indicating if site *s* is a member of fold *c*, and *n*_*s,f*_ is a integer indicating the number of samples from site in the categorical feature class *f*, then we seek to minimize the mean squared error of divergence from perfect stratification:

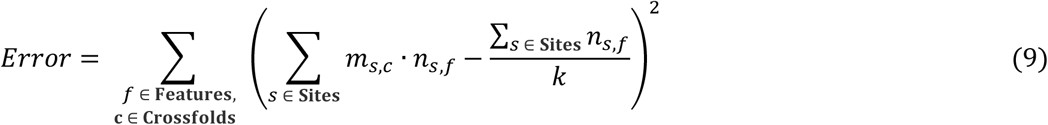

With the constraints that for all sites *s*:

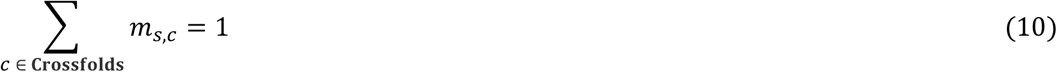

We used CPLEX v12.10, IBM to solve the optimal solution of equations (9) and (10)^69^. Our code used for fold generation for cross-validation is available from https://github.com/fmhoward/PreservedSiteCV. Comparisons between AUROC values and random chance (0.50) was performed using a Z-test with an FDR of 0.05. Comparisons between the false positive rates for African ancestry were performed using a chi-squared test at a tile level.

## Author Contributions

F.M.H. and A.T.P. were responsible for concept proposal and study design. F.M.H., J.D., S.K., and J.N.K. performed essential programming work. J.S., H.C., L.H., and N.C. performed manual oversight and quality control for digital pathology, along with segmentation of tumor. F.M.H., J.D., S.K., D.H., R.N., O.I.O., J.N.K., B.G., and A.T.P. contributed to data interpretation and statistical approaches. All authors contributed to the data analysis and writing of the manuscript.

## Corresponding Authors

Correspondence should be addressed to Alexander T. Pearson or Robert Grossman.

