## Supplemental Tables for "The Impact of Digital Histopathology Batch Effect on Deep Learning Model Accuracy and Bias"

**Supplemental Table 1. Variance in Demographic and Basic Tumor Characteristics by Site, Select Solid Tumors, The Cancer Genome Atlas.**

| Dataset | Variable | # Samples | # Sites | ANOVA p-value |
|---|---|---|---|---|
| **BRCA** | Stage | 998 | 15 | <0.001* |
| | Age | 1016 | 15 | <0.001* |
| | Race | 940 | 15 | <0.001* |
| | Histology | 1017 | 15 | <0.001* |
| | ER | 969 | 15 | 0.002* |
| | PR | 966 | 15 | <0.001* |
| | HER2 | 847 | 15 | 0.002* |
| | BRCA1 | 931 | 15 | 0.009* |
| | Subtype | 914 | 15 | <0.001* |
| | Immune Subtype | 1002 | 15 | <0.001* |
| | 3 Year PFS | 458 | 14 | <0.001* |
| **COAD** | Age | 725 | 9 | <0.001* |
| | Sex | 773 | 9 | <0.001* |
| | Stage | 743 | 9 | <0.001* |
| | Race | 667 | 9 | <0.001* |
| | Histologic Subtype | 760 | 9 | 0.08 |
| | Subsite | 760 | 9 | 0.016* |
| | BMI | 476 | 8 | 0.004* |
| | MSI Status | 416 | 4 | 0.601 |
| | Immune Subtype | 723 | 9 | 0.012* |
| | 3 Year PFS | 336 | 9 | 0.002* |
| **HNSC** | Age | 604 | 10 | <0.001* |
| | Sex | 605 | 10 | <0.001* |
| | Stage | 532 | 9 | <0.001* |
| | Race | 562 | 10 | <0.001* |
| | Grade | 603 | 10 | <0.001* |
| | HPV status | 598 | 10 | <0.001* |
| | Immune Subtype | 598 | 10 | <0.001* |
| | 3 Year PFS | 399 | 9 | <0.001* |
| **KIRC** | Age | 469 | 7 | 0.22 |
| | Sex | 469 | 7 | 0.056 |
| | Stage | 468 | 7 | <0.001* |
| | Race | 411 | 7 | <0.001* |
| | Grade | 469 | 7 | <0.001* |
| | Immune Subtype | 450 | 7 | <0.001* |
| | 3 Year PFS | 334 | 7 | <0.001* |
| **LUAD** | Age | 389 | 8 | 0.682 |
| | Sex | 389 | 8 | 0.002* |
| | Stage | 389 | 8 | <0.001* |
| | Race | 365 | 8 | <0.001* |
| | ALK Translocation Status | 118 | 7 | <0.001* |
| | Immune Subtype | 332 | 8 | <0.001* |
| | 3 Year PFS | 243 | 8 | 0.41 |
| **LUSC** | Age | 340 | 9 | 0.003* |
| | Sex | 348 | 9 | 0.007* |

| | | | |
|---|---|---|---|
| Stage | 348 | 9 | <0.001* |
| Race | 324 | 9 | <0.001* |
| ALK Translocation Status | 157 | 6 | 0.011* |
| Immune Subtype | 343 | 9 | 0.051 |
| 3 Year PFS | 181 | 9 | <0.001* |

(*) indicates p-value remained significant even with Benjamini-Hochberg correction for a false discovery rate of 0.05 (calculated per disease type)

**Supplemental Table 2. ANOVA F-Statistic for First and Second Order Image Features, Select Solid Tumors The Cancer Genome Atlas.**

| Dataset | Statistic | Baseline | Macenko | Reinhard | Grayscale | Grayscale Normalized |
|---|---|---|---|---|---|---|
| BRCA | Red | 40.4* | 13.3* | 15.8* | 78.6* | 23.1* |
| | Red STD | 24.4* | 3.5* | 13.0* | 18.6* | 25.3* |
| | Red Kurtosis | 7.4* | 4.7* | 21.8* | 2.7* | 6.1* |
| | Red Skew | 21.0* | 9.4* | 29.4* | 64.8* | 11.3* |
| | Red Entropy | 23.8* | 23.8* | 23.7* | 23.9* | 23.8* |
| | Green | 73.3* | 37.7* | 12.5* | 78.6* | 23.1* |
| | Green STD | 17.4* | 7.1* | 18.3* | 18.6* | 25.3* |
| | Green Kurtosis | 4.5* | 3.5* | 5.9* | 2.7* | 6.1* |
| | Green Skew | 61.6* | 28.6* | 60.8* | 64.8* | 11.3* |
| | Green Entropy | 24.0* | 23.9* | 23.8* | 23.9* | 23.8* |
| | Blue | 105.5* | 41.7* | 11.0* | 78.6* | 23.1* |
| | Blue STD | 37.2* | 7.3* | 10.5* | 18.6* | 25.3* |
| | Blue Kurtosis | 51.1* | 10.5* | 21.0* | 2.7* | 6.1* |
| | Blue Skew | 123.9* | 37.3* | 29.2* | 64.8* | 11.3* |
| | Blue Entropy | 23.8* | 23.8* | 23.7* | 23.9* | 23.8* |
| | Contrast | 55.9* | 36.8* | 33.2* | 59.5* | 27.2* |
| | Dissimilarity | 41.6* | 33.3* | 31.9* | 44.5* | 26.9* |
| | Homogeneity | 56.8* | 52.0* | 54.7* | 56.8* | 47.4* |
| | ASM | **142.5*** | **152.2*** | **119.5*** | **141.6*** | **141.0*** |
| | Correlation | 66.5* | 43.1* | 42.4* | 68.8* | 46.3* |
| COADREAD | Red | 57.9* | 28.4* | 19.5* | 104.1* | 3.4* |
| | Red STD | 34.9* | 2.2* | 45.3* | 26.1* | 3.5* |
| | Red Kurtosis | 13.9* | 14.9* | 22.6* | 18.9* | 7.5* |
| | Red Skew | 35.6* | 30.5* | 19.2* | 44.4* | 11.2* |
| | Red Entropy | 25.9* | 26.0* | 26.2* | 25.0* | 26.1* |
| | Green | 76.0* | 8.0* | 22.4* | 104.1* | 3.4* |
| | Green STD | 16.9* | 0.7 | 31.1* | 26.1* | 3.5* |
| | Green Kurtosis | 18.3* | 4.6* | 10.6* | 18.9* | 7.5* |
| | Green Skew | 34.5* | 12.3* | 24.7* | 44.4* | 11.2* |
| | Green Entropy | 24.7* | 25.9* | 26.1* | 25.0* | 26.1* |
| | Blue | **232.7*** | 8.6* | 9.2* | 104.1* | 3.4* |
| | Blue STD | 77.4* | 0.9 | 26.7* | 26.1* | 3.5* |
| | Blue Kurtosis | 1.7 | 1.9 | 29.7* | 18.9* | 7.5* |
| | Blue Skew | 125.7* | 13.3* | 32.5* | 44.4* | 11.2* |
| | Blue Entropy | 25.1* | 25.9* | 25.9* | 25.0* | 26.1* |
| | Contrast | 13.3* | 13.9* | 12.7* | 16.3* | 5.8* |
| | Dissimilarity | 28.2* | 27.7* | 28.4* | 31.1* | 12.9* |
| | Homogeneity | 83.4* | 83.2* | 86.2* | 83.3* | 76.1* |
| | ASM | 149.8* | **160.5*** | **149.8*** | **149.3*** | **149.4*** |
| | Correlation | 36.3* | 22.8* | 19.2* | 36.0* | 8.2* |
| HNSC | Red | 19.2* | 12.1* | 12.4* | 23.6* | 1.2 |
| | Red STD | 3.1* | 5.5* | 13.7* | 11.2* | 12.6* |
| | Red Kurtosis | 18.1* | 14.8* | 8.1* | 19.5* | 3.9* |
| | Red Skew | 13.6* | 8.9* | 5.0* | 15.0* | 3.5* |
| | Red Entropy | 1.6 | 1.6 | 1.6 | 1.5 | 1.5 |
| | Green | 31.6* | 17.8* | 19.6* | 23.6* | 1.2 |

| | | | | | | |
|---|---|---|---|---|---|---|
| | Green STD | 9.4* | 11.4* | 13.7* | 11.2* | 12.6* |
| | Green Kurtosis | 17.3* | 13.7* | 5.6* | 19.5* | 3.9* |
| | Green Skew | 16.6* | 3.3* | 7.1* | 15.0* | 3.5* |
| | Green Entropy | 1.4 | 1.5 | 1.6 | 1.5 | 1.5 |
| | Blue | 10.7* | 21.3* | 16.4* | 23.6* | 1.2 |
| | Blue STD | 21.8* | 11.9* | 37.4* | 11.2* | 12.6* |
| | Blue Kurtosis | 22.5* | 4.8* | 9.7* | 19.5* | 3.9* |
| | Blue Skew | 12.1* | 4.1* | 8.7* | 15.0* | 3.5* |
| | Blue Entropy | 1.5 | 1.5 | 1.6 | 1.5 | 1.5 |
| | Contrast | 19.0* | 13.8* | 15.6* | 19.1* | 3.6* |
| | Dissimilarity | 29.4* | 21.9* | 25.2* | 29.2* | 10.1* |
| | Homogeneity | **40.3*** | **40.6*** | **40.7*** | **40.1*** | **39.9*** |
| | ASM | 12.0* | 12.4* | 11.9* | 12.0* | 11.8* |
| | Correlation | 28.1* | 20.7* | 17.1* | 24.4* | 6.9* |
| **LUAD** | Red | 46.4* | 12.9* | 4.7* | 71.1* | 4.0* |
| | Red STD | 45.9* | 5.3* | 21.8* | 33.5* | 4.7* |
| | Red Kurtosis | 10.1* | 3.0* | 12.0* | 3.5* | 7.9* |
| | Red Skew | 24.9* | 10.0* | 11.5* | 31.2* | 29.0* |
| | Red Entropy | 34.1* | 34.8* | 34.8* | 33.4* | 34.8* |
| | Green | 63.4* | 12.6* | 5.9* | 71.1* | 4.0* |
| | Green STD | 28.0* | 4.8* | 3.5* | 33.5* | 4.7* |
| | Green Kurtosis | 3.6* | 4.0* | 4.7* | 3.5* | 7.9* |
| | Green Skew | 21.5* | 17.1* | 25.9* | 31.2* | 29.0* |
| | Green Entropy | 32.2* | 34.8* | 34.8* | 33.4* | 34.8* |
| | Blue | 98.4* | 13.3* | 5.8* | 71.1* | 4.0* |
| | Blue STD | 42.2* | 4.0* | 4.5* | 33.5* | 4.7* |
| | Blue Kurtosis | 17.8* | 4.5* | 41.5* | 3.5* | 7.9* |
| | Blue Skew | 73.0* | 18.5* | 41.7* | 31.2* | 29.0* |
| | Blue Entropy | 34.0* | 34.8* | 34.8* | 33.4* | 34.8* |
| | Contrast | 35.4* | 26.2* | 24.4* | 35.7* | 14.4* |
| | Dissimilarity | 46.3* | 37.2* | 36.6* | 46.9* | 22.0* |
| | Homogeneity | 65.1* | 63.2* | 63.9* | 65.1* | 62.8* |
| | ASM | **126.3*** | **130.1*** | **126.0*** | **126.2*** | **126.2*** |
| | Correlation | 22.6* | 13.2* | 15.2* | 26.0* | 16.6* |
| **LUSC** | Red | 20.1* | 21.2* | 10.9* | **46.9*** | 11.7* |
| | Red STD | 14.8* | 5.1* | 12.5* | 10.6* | 12.1* |
| | Red Kurtosis | 4.6* | 2.1* | 5.6* | 8.9* | 11.4* |
| | Red Skew | 15.7* | 10.9* | 9.6* | 32.0* | 7.5* |
| | Red Entropy | 24.9* | 25.2* | 25.1* | 24.5* | **25.2*** |
| | Green | **51.3*** | 16.8* | 20.1* | **46.9*** | 11.7* |
| | Green STD | 9.8* | 4.6* | 19.5* | 10.6* | 12.1* |
| | Green Kurtosis | 10.3* | 8.0* | 10.3* | 8.9* | 11.4* |
| | Green Skew | 32.8* | 20.4* | **29.6*** | 32.0* | 7.5* |
| | Green Entropy | 23.8* | **25.3*** | 25.2* | 24.5* | **25.2*** |
| | Blue | 45.6* | 17.0* | 18.5* | **46.9*** | 11.7* |
| | Blue STD | 14.3* | 4.2* | 17.8* | 10.6* | 12.1* |
| | Blue Kurtosis | 4.5* | 2.9* | 7.5* | 8.9* | 11.4* |
| | Blue Skew | 27.5* | 16.9* | 12.9* | 32.0* | 7.5* |
| | Blue Entropy | 24.8* | 25.2* | 25.2* | 24.5* | **25.2*** |

| | | | | | | |
|---|---|---|---|---|---|---|
| | Contrast | 14.9* | 10.7* | 12.2* | 15.0* | 5.8* |
| | Dissimilarity | 14.1* | 9.0* | 10.4* | 14.3* | 4.4* |
| | Homogeneity | 11.9* | 11.5* | 11.6* | 12.0* | 11.0* |
| | ASM | 9.7* | 10.7* | 8.5* | 9.7* | 9.6* |
| | Correlation | 21.5* | 15.5* | 16.4* | 22.8* | 13.3* |
| **KIRC** | Red | 15.5* | 21.5* | 1.8 | 25.5* | 16.6* |
| | Red STD | 23.6* | 15.4* | 4.4* | 42.1* | 16.0* |
| | Red Kurtosis | 19.6* | 6.2* | 19.4* | 13.8* | 12.0* |
| | Red Skew | 20.3* | 14.7* | 22.3* | 11.4* | 7.0* |
| | Red Entropy | 27.3* | 27.8* | 27.8* | 26.3* | 28.0* |
| | Green | 24.0* | 11.8* | 1.6 | 25.5* | 16.6* |
| | Green STD | 39.1* | 17.5* | 12.9* | 42.1* | 16.0* |
| | Green Kurtosis | 11.4* | 7.7* | 14.7* | 13.8* | 12.0* |
| | Green Skew | 11.3* | 9.3* | 11.2* | 11.4* | 7.0* |
| | Green Entropy | 25.5* | 28.1* | 27.9* | 26.3* | 28.0* |
| | Blue | 30.7* | 9.7* | 3.9* | 25.5* | 16.6* |
| | Blue STD | 49.8* | 16.9* | 11.2* | 42.1* | 16.0* |
| | Blue Kurtosis | 13.6* | 8.5* | 19.7* | 13.8* | 12.0* |
| | Blue Skew | 8.4* | 6.8* | 11.8* | 11.4* | 7.0* |
| | Blue Entropy | 26.6* | 27.9* | 27.8* | 26.3* | 28.0* |
| | Contrast | 36.1* | 14.6* | 15.9* | 39.1* | 8.3* |
| | Dissimilarity | 40.8* | 20.9* | 23.5* | 42.6* | 11.1* |
| | Homogeneity | 61.7* | 59.0* | 59.2* | 61.8* | 57.1* |
| | ASM | **102.2*** | **102.5*** | **96.7*** | **102.1*** | **101.2*** |
| | Correlation | 10.7* | 13.2* | 8.0* | 9.6* | 7.9* |

(*) indicates p-value remained significant even with Benjamini-Hochberg correction for a false discovery rate of 0.05 (calculated per disease type, per stain normalization method). Factor with highest associated F-statistic indicated in bold.

**Supplemental Table 3. One-Versus-Rest Area under the Receiver Operating Characteristic Curve (AUROC) for Prediction of Tissue Submitting Site, with 3-Fold Cross Validation.**

| Dataset | Slide Adjustment Method | CV1 AUROC | CV2 AUROC | CV3 AUROC | Average | T-Test p-value Compared to Baseline |
|---|---|---|---|---|---|---|
| **BRCA** | Baseline | 0.972 | 0.985 | 0.985 | 0.981 | --- |
| | Macenko | 0.968 | 0.97 | 0.978 | 0.972 | 0.119 |
| | Reinhard | 0.985 | 0.981 | 0.981 | 0.982 | 0.796 |
| | Grayscale | 0.948 | 0.946 | 0.944 | 0.946 | 0.023* |
| | Grayscale Normalized | 0.957 | 0.942 | 0.95 | 0.95 | 0.065 |
| | Light HSV Augmentation | 0.958 | 0.967 | 0.966 | 0.964 | 0.008* |
| | Heavy HSV Augmentation | 0.953 | 0.956 | 0.967 | 0.957 | 0.025* |
| **COADREAD** | Baseline | 0.985 | 0.984 | 0.988 | 0.986 | --- |
| | Macenko | 0.968 | 0.982 | 0.978 | 0.976 | 0.155 |
| | Reinhard | 0.957 | 0.978 | 0.983 | 0.973 | 0.225 |
| | Grayscale | 0.950 | 0.968 | 0.966 | 0.961 | 0.049 |
| | Grayscale Normalized | 0.930 | 0.969 | 0.963 | 0.954 | 0.119 |
| | Light HSV Augmentation | 0.961 | 0.969 | 0.974 | 0.968 | 0.031 |
| | Heavy HSV Augmentation | 0.955 | 0.971 | 0.976 | 0.967 | 0.088 |
| **HNSC** | Baseline | 0.984 | 0.934 | 0.981 | 0.966 | --- |
| | Macenko | 0.935 | 0.944 | 0.935 | 0.938 | 0.278 |
| | Reinhard | 0.924 | 0.913 | 0.921 | 0.919 | 0.069 |
| | Grayscale | 0.899 | 0.835 | 0.834 | 0.856 | 0.028 |
| | Grayscale Normalized | 0.886 | 0.849 | 0.870 | 0.868 | 0.006* |
| | Light HSV Augmentation | 0.959 | 0.891 | 0.929 | 0.926 | 0.037 |
| | Heavy HSV Augmentation | 0.941 | 0.871 | 0.897 | 0.903 | 0.033 |
| **LUAD** | Baseline | 0.979 | 0.990 | 0.955 | 0.975 | --- |
| | Macenko | 0.952 | 0.977 | 0.937 | 0.955 | 0.042 |
| | Reinhard | 0.960 | 0.976 | 0.933 | 0.956 | 0.016* |
| | Grayscale | 0.890 | 0.931 | 0.871 | 0.897 | 0.014* |
| | Grayscale Normalized | 0.914 | 0.934 | 0.892 | 0.913 | 0.002* |
| | Light HSV Augmentation | 0.928 | 0.957 | 0.934 | 0.940 | 0.057 |
| | Heavy HSV Augmentation | 0.916 | 0.952 | 0.910 | 0.926 | 0.023* |
| **LUSC** | Baseline | 0.984 | 0.985 | 0.990 | 0.986 | --- |
| | Macenko | 0.957 | 0.944 | 0.979 | 0.960 | 0.093 |
| | Reinhard | 0.963 | 0.967 | 0.977 | 0.969 | 0.018* |
| | Grayscale | 0.939 | 0.916 | 0.945 | 0.933 | 0.022* |
| | Grayscale Normalized | 0.939 | 0.922 | 0.953 | 0.938 | 0.024* |
| | Light HSV Augmentation | 0.947 | 0.956 | 0.971 | 0.958 | 0.032* |
| | Heavy HSV Augmentation | 0.941 | 0.929 | 0.953 | 0.941 | 0.015* |
| **KIRC** | Baseline | 0.996 | 0.999 | 0.999 | 0.998 | --- |
| | Macenko | 0.992 | 0.991 | 0.994 | 0.992 | 0.042 |
| | Reinhard | 0.990 | 0.995 | 0.990 | 0.992 | 0.049 |
| | Grayscale | 0.950 | 0.883 | 0.940 | 0.924 | 0.076 |
| | Grayscale Normalized | 0.956 | 0.949 | 0.948 | 0.951 | 0.006* |
| | Light HSV Augmentation | 0.974 | 0.978 | 0.966 | 0.973 | 0.022 |
| | Heavy HSV Augmentation | 0.960 | 0.971 | 0.949 | 0.960 | 0.027 |

(*) indicates p-value remained significant even with Benjamini-Hochberg correction for a false discovery rate of 0.05, calculated per disease type.

**Supplemental Table 4. One-Versus-Rest Area under the Receiver Operating Characteristic Curve (AUROC) for Prediction of Genomic Ancestry, with 3-Fold Cross Validation.**

| Slide Adjustment Method | Average (95% CI) Balanced by Race | P-Value | Average (STD) Sites Preserved | P-Value |
|---|---|---|---|---|
| Baseline | 0.784 (0.747 - 0.821) | < 0.001* | 0.562 (0.502 - 0.622) | 0.022 |
| Macenko | 0.760 (0.722 - 0.798) | < 0.001* | 0.522 (0.423 - 0.621) | 0.334 |
| Reinhard | 0.772 (0.729 - 0.815) | < 0.001* | 0.589 (0.472 - 0.707) | 0.068 |
| Grayscale | 0.687 (0.626 - 0.748) | < 0.001* | 0.566 (0.472 - 0.661) | 0.084 |
| Grayscale Normalized | 0.727 (0.678 - 0.775) | < 0.001* | 0.575 (0.457 - 0.693) | 0.105 |
| Light HSV Augmentation | 0.759 (0.746 - 0.772) | < 0.001* | 0.552 (0.422 - 0.682) | 0.217 |
| Heavy HSV Augmentation | 0.732 (0.726 - 0.738) | < 0.001* | 0.559 (0.445 - 0.673) | 0.156 |

P-values listed for Z-test comparison to an AUROC of 0.5, with significant indicating a better than random performance at predicting race. Significant p-values corrected for a false discovery rate of 0.05 denoted with *.

**Supplemental Table 5. False Positive Rate for Prediction of European Ancestry, Balanced by Race versus Preserved Sites.**

| Slide Adjustment Method | False Positive AFR Balanced by Race (%) | False Positive AFR Preserved Sites (%) | Chi-Squared p-value |
|---|---|---|---|
| Baseline | 80.6 | 0.1 | < 0.001* |
| Macenko | 37.4 | 3.4 | < 0.001* |
| Reinhard | 48.1 | 3.4 | < 0.001* |
| Grayscale | 36.7 | 14.1 | < 0.001* |
| Grayscale + Normalization | 38.8 | 9.3 | < 0.001* |
| Light HSV Augmentation | 49.7 | 6.0 | < 0.001* |
| Heavy HSV Augmentation | 67.8 | 14.6 | < 0.001* |

False positive rates are calculated at a tile level for patients enrolled from University of Chicago, the site with the highest proportion of African ancestry. A high rate of false positive predictions of African Ancestry are seen when the site of interest is split into both the validation and testing sets, suggestive that the model is learning that patients with the staining / slide image patterns characteristic of University of Chicago are more likely to be African. However, when models are trained from other institutions, false positive African ancestry is dramatically reduced. Slide normalization techniques improve the false positive rate, but not to the same level as isolating sites to either validation or testing sets. (*) indicates p-value remained significant even with Benjamini-Hochberg correction for a false discovery rate of 0.05
